# Engineering serine metabolism to enhance AOX1 promoter self-induction in a formate dehydrogenase-deficient *Komagataella phaffii*

**DOI:** 10.1101/2025.07.04.663172

**Authors:** Rocio Cozmar, Julio Berrios, Patrick Fickers

## Abstract

The methylotrophic yeast *Komagataella phaffii* is a premier host for recombinant protein (rProt) production, traditionally relying on methanol induction of the alcohol oxidase 1 promoter (pAOX1). However, methanol flammability and associated industrial limitations have motivated the search for methanol-free induction systems. We recently demonstrated that disruption of formate dehydrogenase (FDH) in *K. phaffii* allows endogenous formate, derived from tetrahydrofolate (THF)-mediated C1 metabolism, to induce pAOX1 without addition of external inducers. Building on this, we hypothesized that increasing intracellular formate production by enhancing serine biosynthesis could further improve promoter induction and rProt productivity. Overexpression of SER3, encoding 3-phosphoglycerate dehydrogenase, the rate-limiting enzyme in serine synthesis, significantly increased pAOX1-driven expression of an intracellular reporter protein (eGFP) and secreted glucose oxidase (Gox) from *Aspergillus niger*, without compromising cell fitness. Enhanced formate accumulation and stronger pAOX1 induction were observed in both micro- and bioreactor cultivations using sorbitol or glycerol-sorbitol mixtures. In bioprocess conditions, SER3 overexpression led to a 30% increase in specific Gox activity compared to the parental FdhKO strain. This study provides a cost-effective metabolic engineering strategy for methanol-free, self-inducible expression systems in *K. phaffii* based on pAOX1, enabling safer and more sustainable industrial rProt production.

## Introduction

The methylotrophic yeast *Komagataella phaffii* (formerly *Pichia pastoris*) is widely used for recombinant protein (rProt) production, primarily relying on expression systems based on the tightly regulated promoter of the alcohol oxidase 1 gene (pAOX1). This promoter is repressed in the presence of glucose or glycerol and strongly induced by methanol. However, methanol use at an industrial scale poses several challenges, including flammability, high oxygen demand during catabolism, and stringent safety regulations (Cereghino & Cregg, 2000; Potvin et al., 2012). These limitations have spurred the development of methanol-free induction strategies, including novel regulatory systems or alternative carbon sources.

Formate has emerged as a promising alternative inducer for pAOX1 (Jayachandran et al., 2017; Singh & Narang, 2020; Tyurin & Kozlov, 2015). As a C1 molecule with low energy content, formate must be supplemented along with a non-repressive carbon source during the rProt production phase. Sorbitol has proven to be an effective co-substrate in this context (Niu et al., 2013; Singh & Narang, 2023). Metabolically, formate is an intermediate in the methanol dissimilative pathway, where it is oxidized to carbon dioxide by formate dehydrogenase (FDH) (Bustos et al., 2022). Additionally, formate is a key intermediate in the tetrahydrofolate-mediated one-carbon (THF-C1) metabolism, which contributes to several anabolic processes, including de novo purine synthesis (Christensen & MacKenzie, 2006; Supplementary information Figure S1). The THF-C1 pathway operates in both the mitochondria and cytoplasm, with formate serving as a C1 shuttle between the two compartments.

Recently, we demonstrated that *K. phaffii* cells deficient in FDH (FdhKO), when cultivated on sorbitol-based medium, accumulate formate endogenously through THF-C1 metabolism. This intracellular formate was sufficient to trigger pAOX1 induction without any external inducer. This finding is particularly significant, as it enables the conversion of any pAOX1-based expression system into a self-inducing system in an FdhKO background. When such cells are grown on a glycerol-sorbitol mixture, rProt synthesis initiates at the end of the exponential growth phase, upon glycerol depletion from the medium (Bustos et al., 2024)

In THF-C1 metabolism, formate is derived from serine. We previously showed that serine supplementation markedly increases pAOX1 induction and rProt productivity (Bustos et al., 2024). Serine is synthesized from the glycolytic intermediate 3-phosphoglycerate through three enzymatic steps involving phosphoglycerate dehydrogenase (Pgdh), phosphoserine aminotransferase (Psat1), and phosphoserine phosphatase (Psph), encoded by SER3, SER1, and SER2, respectively (Kobayashi et al., 2022; Pruneski et al., 2011, Figure 1). In *Saccharomyces cerevisiae*, this pathway is tightly regulated as serine represses SER1 transcription (Delic et al., 2013), while SER3 expression is modulated by the non-coding RNAi SRG1 (Martens et al., 2005; Pruneski et al., 2011) In addition, serine can be reversibly converted from glycine by serine hydroxymethyltransferases Shm1 and Shm2, which also participate in THF-C1 metabolism (Esch et al., 2020; Supplementary information Figure S1).

**Figure 1.**
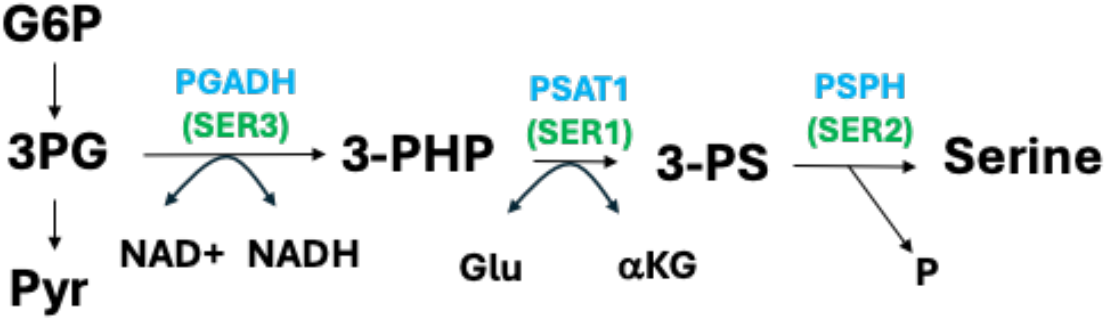
Serine biosynthesis pathway in *K. phaffii*. Abbreviation are as follows: PGADH, 3- phophoglicerate dehydrogenase; PSAT1, phosphoserine aminotransferase encoded; PSPH, phosphoserine phosphatase; G6P, glucose-6-phosphate; 3PG, 3-phosphoglycerate; NAD+, nicotinamide adenine dinucleotide oxidized; NADH, nicotinamide adenine dinucleotide reduced; Glu, glutamate; αKG, alfa-ketoglutarate; Pyr, pyruvate; P, phosphate.

As serine supplementation is not economically viable at an industrial scale, we further engineered the FdhKO strain by overexpressing the SER genes individually or in combination. This strategy was designed to enhance intracellular formate accumulation and thereby strengthen AOX1-driven gene expression under methanol-free conditions. Specifically, overexpression of SER3 led to an increased formate accumulation and, consequently, higher pAOX1 induction. This effect was exemplified by a 30% increase in the specific activity of secreted glucose oxidase from *Aspergillus niger*, used here as a model heterologous rProt.

## Materials and methods

### Strains and media and culture conditions

The *Escherichia coli* and *K. phaffii* strains used in this study are listed in Table 1 and supporting information Table S1. *E. coli* was grown at 37°C in Luria-Bertani medium (LB), supplemented with antibiotics as follows: 100 µg mL^-1^ ampicillin, 50 µg mL^-1^ kanamycin, 25 µg mL^-1^ zeocin, or 50 µg mL^-1^ nourseothricin. *K. phaffii* was grown at 30 °C either on YPD medium (20 g L^-1^ glucose, 10 g L^-1^ Difco yeast extract, and 10g L^-1^ Difco bacto peptone) or in YNB medium (1.7 g L^-1^ Difco YNB w/o ammonium chloride and amino acids, 5 g L^-1^ NH_4_Cl and, 0.4 mg L^-1^ biotin, 100 mM potassium phosphate buffer, pH 6.0) supplemented with 1.75 g L^-1^ glycerol and 3.5 g L^-1^ sorbitol (YNBGS4), 20 g L^-1^ sorbitol (YNBS) or 20 g L^-1^ glycerol and 20 g L^-1^ sorbitol (YNBGS). *K. Phaffii* transformants were selected on YPD agar plates, supplemented with 25 µg mL^-1^ zeocin (YPD-Zeo), 200 µg mL^-1^ Hygromycin (YPD-Hyg), 500 µg mL^-1^ geneticin (YPD-Genet) or 100 µg mL^-1^ nourseothricin (YPD-NAT). Precultures were operated as previously described (Bustos et al., 2024). Shake flask cultures were performed either in 250 ml Erlenmeyer containing 25 ml of medium with agitation of 150 rpm. Cultures in microbioreactors (BioLector 2, m2p-labs, Baesweiler, Germany), were operated at 1000 rpm with relative humidity of 85 % using 48-well Flower plates (M2P-MTP-48B, Beckman Coulter) containing 1 ml of medium. Stirred tank bioreactor cultivations were performed in 155 ml medium in 200-ml DASGIP bioreactors (DASGIP DASbox Mini Bioreactors SR0250ODLS, Eppendorf, Hamburg, Germany at 30°C) as described previously (Theron et al., 2019). Operating parameters were as follows: aeration at 0.5 vvm, stirring at 900 rpm, and pH regulated at pH 6.

**TABLE 1.** *K. phaffii* strains used in this study.

| Number | Names | Parental strain, relevant genotype | Reference |
| --- | --- | --- | --- |
| RIY232 | GS115pro | GS115, HIS4 | (Theron et al., 2019) |
| RIY230 | Fdh eGfp | GS115, pAOX1-eGFP, Zeo<sup>+</sup> | (Bustos et al., 2024) |
| RIY536 | FdhKO eGfp Z<sup>+</sup> | RIY230, fdh1Δ, pAOX1-eGFP, Zeo<sup>+</sup> | (Bustos et al., 2024) |
| RIY540 | eGfp | RIY536, fdh1Δ, pAOX1-eGFP, Zeo<sup>-</sup> | (Bustos et al., 2024) |
| RIY657 | gSER1-eGFP | RIY540, fdh1Δ, pGAP-SER1, pAOX1-eGFP, Zeo<sup>+</sup> | This work |
| RIY658 | mSER1-eGFP | RIY540, fdh1Δ, pPMDH3-SER1, pAOX1-eGFP, Zeo<sup>+</sup> | This work |
| RIY659 | gSER2-eGFP | RIY540, fdh1Δ, pGAP-SER2, pAOX1-eGFP, Zeo<sup>+</sup> | This work |
| RIY660 | *mSER2*-eGFP | RIY540, *fdh1Δ*, pPMDH3-SER2, pAOX1-eGFP, Zeo<sup>+</sup> | This work |
| RIY661 | *gSER3*-eGFP | RIY540 *fdh1Δ*, pGAP-SER3, pAOX1-eGFP, Zeo<sup>+</sup> | This work |
| RIY662 | *mSER3*-eGFP | RIY540, *fdh1Δ*, pPMDH3-SER3, pAOX1-eGFP, Zeo<sup>+</sup> | This work |
| RIY663 | *mSER3-gSER1*-eGFP | RIY662, *fdh1Δ*, pPMDH3-SER3, pGAP-SER1, pAOX1-eGFP, Zeo<sup>+</sup>, Hyg<sup>+</sup> | This work |
| RIY664 | *mSER3-mSER1*-eGFP | RIY662, *fdh1Δ*, pPMDH3-SER3, pPMDH3-SER1, pAOX1-eGFP, Zeo<sup>+</sup>, Hyg<sup>+</sup> | This work |
| RIY538 | FdhKO Z<sup>+</sup> | RIY232, *fdh1Δ*, Zeo<sup>+</sup> | This work |
| RIY539 | FdhKO | RIY538, *fdh1Δ* | This work |
| RIY655 | GOX | RIY539, *fdh1Δ*, pAOX1-αMF-GOX, Zeo<sup>+</sup> | This work |
| RIY665 | GOX-SER3 | RIY655, *fdh1Δ*, pAOX1-αMF-GOX, pMDH3-SER3, Zeo<sup>+</sup>, NAT<sup>+</sup> | This work |

### Construction of plasmids and *K. phaffii* strains

The general genetic techniques were as described elsewhere (Bustos et al., 2024). The plasmids and primers used are listed in Supporting information Table S1 and S2. The DNA sequence of signal sequence of the alfa mating factor (*α*MF) from *Saccharomyces cerevisiae* (NM_001184001.1) and the intron less coding sequence of the glucose oxidase (Gox) gene from *A. niger* (x60601.1) were codon optimized for *K. phaffii* using SnapGene software. Codons with a frequency below 10% were replaced with the most abundant one coding for the same amino acid. Recognition sequence for *Bsa*I or *Bpi*I were added as shown in Supporting information Figures S1 and S2. The *α*MF and Gox synthetic DNA fragments obtained from IDT technologies (Leuven Belgium) were cloned into pUC57 and pGEMT-easy vector, respectively, to yield plasmids RIE457 and RIE585, respectively (Supporting information Table S1). The *α*MF (RIE457) was digested by *Bsa*I and ligated at the corresponding site of plasmid A2 (BB1_23), yielding plasmids RIE504 (A2_BB1_23_*αMFGOX*). Plasmid RIE507 (BB3aZ_14-p*AOX1*-*αMF- GOX*-*SsCy1*tt) was obtained by GoldenGate assembly from plasmids B8 (BB1_12_p*AOX1*), C1 (BB1_34_ScCYC1tt), D12 (BB3aZ_14), with RIE504, using *Bpi*I as the restriction enzyme as describe elsewhere (Prielhofer et al., 2017). The correctness of the assembly was verified by DNA sequencing using primers pair CkBB3aZ14-Fw/CkBB3aZ1-Rv.

The genes PAS_chr3_0566 (SER1), PAS_chr4_0285 (SER2), and PAS_chr2-1_0657 (SER3) were PCR-amplified from *K. Phaffii* GS115pro genomic DNA used as a template using primer pairs SER1-Fw/SER1- Rv; SER2-Fw/SER2-Rv and SER3-Fw/SER3-Rv, respectively. The resulting PCR fragments were cloned into Zero Blunt TOPO vector, yielding plasmids RIE508 (TOPO_*SER1*), RIE509 (TOPO_*SER2*), and RIE510 (TOPO_*SER3*), respectively. Positive constructs were verified by DNA sequencing with M13-Fw/M13- Rv primers. To construct the SER gene expression vectors, promoters (pGAP, pMDH3), SER genes and terminator (ScCYC1tt) were first PCR amplified using specific primer pairs designed to generate overlap that allows subsequent assembly of the expression vector using the NEBuilder Hifi DNA assembly, NewEngland Biolabs. The pGAP promoter fragments were PCR amplified from plasmid A4 (BB1_12_p*GAP*) using primer pairs 1.pGAP.SER1-Rv/2.pGAP-Fw; 1.pGAP.SER2-Rv/2.pGAP-Fw; 1.pGAP.SER3-Rv/2.pGAP-Fw to generate parts 1, 2 and 3, respectively. The pMDH3 promoter fragments were PCR amplified from plasmid A9 (BB1_12_P*MDH3*) using primer pairs 1.pMDH3.SER1- Rv/2.pMDH3-Fw; 1.pMDH3.SER2-Rv/2.pMDH3-Fw, 1.pMDH3.SER3-Rv/2.pMDH3-Fw, 1.pMDH3.SER3- Rv/1.BB3rN14pMDH3-Fw to generate parts 4, 5, 6 and 7, respectively. The SER1 gene fragments were amplified from plasmid RIE508 (TOPO_*SER1*) using primer pairs 3.SER1Rv/2.pGAP.SER1Fw, 3.SER1Rv/2.pMDH3.SER1Fw to generate parts 8 and 9, respectively. The SER2 gene fragment were amplified from plasmid RIE509 (TOPO_*SER2*) using primer pairs 3.SER2Rv/2.pGAP.SER2Fw, 3.SER2Rv/2.pMDH3.SER2Fw to generate parts 10 and 11, respectively. The SER3 gene fragments were amplified from plasmid RIE510 (TOPO_*SER3*) using primer pairs 3.SER3Rv/2.pGAP.SER3Fw, 3.SER3Rv/2.pMDH3.SER3Fw to generate parts 12 and 13, respectively. The ScCYC1 terminator fragments were PCR amplified from plasmid C1 (BB1_34_ScCYC1tt) using primer pairs 4.CYC1- Rv/4.SER1.C1-Fw; 4.BB3eH14.C1-Rv/4.SER1.C1-Fw, 4.CYC1-Rv/4.SER2.C1-Fw; 4.CYC1-Rv/4.SER3.C1- Fw, 4.BB3rN14.C1-Rv/4.SER3.C1-Fw to generate parts 14, 15, 16, 17 and 18 respectively. All the obtained PCR fragments were gel purified and quantified with a nanodrop (Thermo Fisher Scienfic). The final expression vector were obtained by assembly of parts as follows : parts 1, 8, 14 and D12 plasmid *Bpi*I digested to obtain plasmid RIP511 (BB3aZ_14-p*GAP*-*SER1*-*SsCy1*tt); parts 4, 9, 14 and D12 plasmid *Bpi*I digested to obtain plasmid RIP512 (BB3aZ_14-p*MDH3*-*SER1*-*SsCy1*tt); parts 2, 10, 16 and D12 plasmid *Bpi*I digested to obtain plasmid RIP513 (BB3aZ_14-p*GAP*-*SER2*-*SsCy1*tt); parts 5, 11, 16 and D12 plasmid *Bpi*I digested to obtain plasmid RIP514 (BB3aZ_14-p*MDH3*-*SER2*-*SsCy1*tt); parts 1, 12, 17 and D12 plasmid *Bpi*I digested to obtain plasmid RIP515; parts 1, 12, 17 and D12 plasmid *Bpi*I digested to obtain plasmid RIP515; parts 6, 13, 17 and D12 plasmid *Bpi*I digested to obtain plasmid RIP516; parts 1, 8, 15 and E1 plasmid *Bpi*I digested to obtain plasmid RIP517; parts 4, 9, 15 and E1 plasmid *Bpi*I digested to obtain plasmid RIP518; parts 7, 4, 18 and E8 plasmid *Bpi*I digested to obtain plasmid RIP519. The correctness of the assembly constructs was verified by PCR with primers pair CkBB3aZ14-Fw/CkBB3aZ1-Rv for RIP511, RIP512, RIP513, RIP, RIP514, RIP515 and RIP516; 1BB3aZ14- Rv/4.BB3eH14-Fw for RIP517, RIP518 and 1.BB3rN14-Rv/4.BB3rN14-Fw for RIP519. Correctness of the plasmid sequence was verified by DNA sequencing.

The construction of yeast strains RIY230 (Fdh eGfp), RIY536 (FdhKO eGfp Z^+^), RIY540 (eGfp) is described elsewhere (Bustos et al., 2024). Strains RIY657 (*fdh1Δ*, p*GAP-SER1*, p*AOX1-eGFP*, hereafter *g*SER1-eGfp), RIY658 (*fdh1Δ*, p*MDH3-SER1*, p*AOX1-eGFP*, hereafter *m*SER1-eGFP), RIY659 (*fdh1Δ*, p*GAP-SER2*, p*AOX1-eGFP*, hereafter *g*SER2-eGFP), RIY660 (*fdh1Δ*, p*MDH3-SER2*, p*AOX1-eGFP*, hereafter mSER2-eGFP strain), RIY661 (*fdh1Δ*, p*GAP-SE31*, p*AOX1-eGFP*, hereafter *g*SER3-eGFP), RIY662 (*fdh1Δ*, p*MDH3-SER3*, p*AOX1-eGFP*, hereafter *m*SER3-eGFP strain) were obtained by transformation of strain RIY540 with *Asc*I linearized plasmids RIP511, RIP512, RIP513, RIP, RIP514, RIP515, RIP516, respectively. Strains RIY663 (*fdh1Δ*, pPMDH3*-SER3*, pGAP*-SER1*, p*AOX1-eGFP*, hereafter *m*SER3-*g*SER1-eGFP strain) and RIY664 (*fdh1Δ*, pPMDH3*-SER3*, pMDH3*-SER1*, p*AOX1-eGFP, m*SER3-*m*SER1-eGFP) were obtained by transformation of strain RIY662 with *Pme*I linearized plasmids RIP517 and RIP518 respectively.

For marker rescue, RIY538 (FdhKO Z^+^) strain was transformed with the replicative vector RIP396 (pKTAC-Cre) and transformants were selected on YPD-Genet. The resulting strain was RIY539 (FdhKO). RIY655 (*fdh1Δ*, p*AOX1*-*αMF-GOX*, hereafter GOX) strain was obtained by transformation of strain RIY539 with *Asc*I linearized plasmids RIP507 while RIY665 strain (*fdh1Δ*, p*AOX1*-*αMF-GOX, pMDH3- SER3*, hereafter SER3-GOX3 strain) was obtained by transformation of strain RIY655 with *Asc*I linearized plasmids RIP519. Correctness of the genotype of RIY655, RIY657, RIY658, RIY659, RIY660, RIY661 and RIY662 strains were verified by PCR using the primers CkBB3aZ14-Fw/CkBB3aZ1-Rv while that of RIY663 and RIY664 strains was made using the primers 1BB3aZ14-Rv/4.BB3eH14-Fw. The genotype of the RIY665 strain was verified by PCR using the primers 1.BB3rN14-Rv/4.BB3rN14-Fw. Each primer pairs annealed to the at the 5′ and 3′ regions of the integrated constructed. A schematic representation of the strain genotype is presented in Supplementary information Figure S4.

### Analytical methods

The cell growth was monitored either by optical density at 600nm (OD600) or dry cell weight (DCW) as previously described (Carly et al., 2016). Sorbitol and glycerol concentrations were determined using high-performance liquid chromatography (Agilent 1100 series equipped with UV (210 nm) and RID detector (Agilent Technologies, Santa Clara, CA, USA) using an Aminex HPX-87H column (300 × 7.8 mm Bio-Rad, Hercules, CA, USA). Compounds were eluted from the column at 65 °C with a flow rate of 0.5 mL min^−1^ and using a 5 mM H_2_SO_4_ solution as the mobile phase. During cultures in microbioreactors biomass was monitored by light scattering at 600 nm (gain set at 2) while fluorescence was quantified at 520 nm (excitation 488 nm, gain set at 4) every 10 min. The eGFP fluorescence was expressed as specific fluorescence (sFU). For that purpose, fluorescence values were divided by the biomass values. The specific fluorescence value of the parental strain GS115pro (RIY232) was subtracted from the specific fluorescence of eGFP-producing strains to normalize the data.

Formate concentration in the culture supernatant were measured using the formic acid assay kit (Megazyme Inc., Bray, Ireland) according to the manufacturer instructions. Briefly, formate is converted into carbon dioxide by formate dehydrogenase with the formation of NADPH that is quantified at 340 nm. Formate concentration is deduced from a calibration curve.

Glucose oxidase activity was monitored using a glucose oxidase assay kit (Megazyme Inc., Bray, Ireland) according to the manufacturer instructions. Briefly, Gox catalyzes the oxidation of β-D-glucose to D-glucono-δ-lactone with the concurrent release of hydrogen peroxide. In the presence of peroxidase, the H_2_O_2_ generated is converted, in the presence of p-hydroxybenzoic acid, into quinoneimine which is measured at 510 nm. Gox activity is calculated based on a standard curve. For the formate and enzymatic assay, absorbances at specific wavelength were monitored over time using a SpectraMax M2 (Molecular Devices, San Jose, CA, USA). All assays were conducted in triplicates.

## Results and discussion

### Overexpression of the 3-phosphoglycerate dehydrogenase-encoding gene in a FdhKO strain increases formate accumulation and pAOX1 induction

Serine is a metabolic precursor in various anabolic pathways and serves as a donor of one-carbon units in the THF-C1 pathway, where formate is a key metabolite. We previously demonstrated that endogenous formate induces the pAOX1 promoter in an FdhKO strain, and that supplementing the culture medium with serine positively influences pAOX1 induction and, consequently, rProt productivity (Bustos et al., 2024).

We hypothesized that enhancing serine biosynthesis could increase intracellular formate levels in a FdhKO strain and thereby further potentiate pAOX1 induction. To test this, genes involved in serine synthesis from 3-phosphoglycerate, namely SER1 (Sat1), SER2 (Psph), and SER3 (Pgadh) (Supplementary information Table S3), were overexpressed in a FdhKO strain expressing eGFP under the control of the pAOX1 promoter (fdh1Δ, pAOX1-eGFP, hereafter eGFP strains). All the strains used in this study are disrupted for FDH (FdhKO strains), however for simplicity this was not mentioned in gene name nomenclature (Table 1). Each gene was expressed under the control of either the strong pGAP (glyceraldehyde-3-phosphate dehydrogenase) promoter or the weak pMDH3 (malate dehydrogenase 3) promoter. The resulting strains were designated as follows: *g*SER1-eGFP (fdh1Δ, pAOX1-eGFP, pGAP-SER1), *m*SER1-eGFP (fdh1Δ, pAOX1-eGFP, pMDH3-SER1), *g*SER2-eGFP (fdh1Δ, pAOX1-eGFP, pGAP-SER2), *m*SER2-eGFP (fdh1Δ, pAOX1-eGFP, pMDH3-SER2), *g*SER3-eGFP (fdh1Δ, pAOX1-eGFP, pGAP-SER3), *m*SER3-eGFP (fdh1Δ, pAOX1-eGFP, pMDH3-SER3) (Table 1). These strains were cultivated in microbioreactors in sorbitol medium (YNBS), and specific eGFP fluorescence (i.e. normalized to biomass) was monitored as a proxy to estimate pAOX1 induction every 10 minutes over 48 h. For strains overexpressing SER1 (Sat1) and SER2 (Psph), the specific fluorescence was lower than that of the parental eGFP strain, regardless of the promoter strength used (Figure 2A, B). In contrast, overexpression of SER3 encoding 3-phosphoglycerate dehydrogenase (Pgadh) resulted in a marked increase in specific fluorescence and thus pAOX1 induction, particularly when expressed under the pMDH3 promoter (1.28-fold vs. 1.18-fold for pGAP after 48 h; Figure 2C). Pgadh catalyses the first reaction of the serine biosynthesis pathway, which has been identified as the rate-limiting step of serine synthesis in *S. cerevisiae* (Esch et al., 2020). Based on our observation, this step is also the rate limiting for serine anabolism in *K. phaffii*.

**Figure 2:**
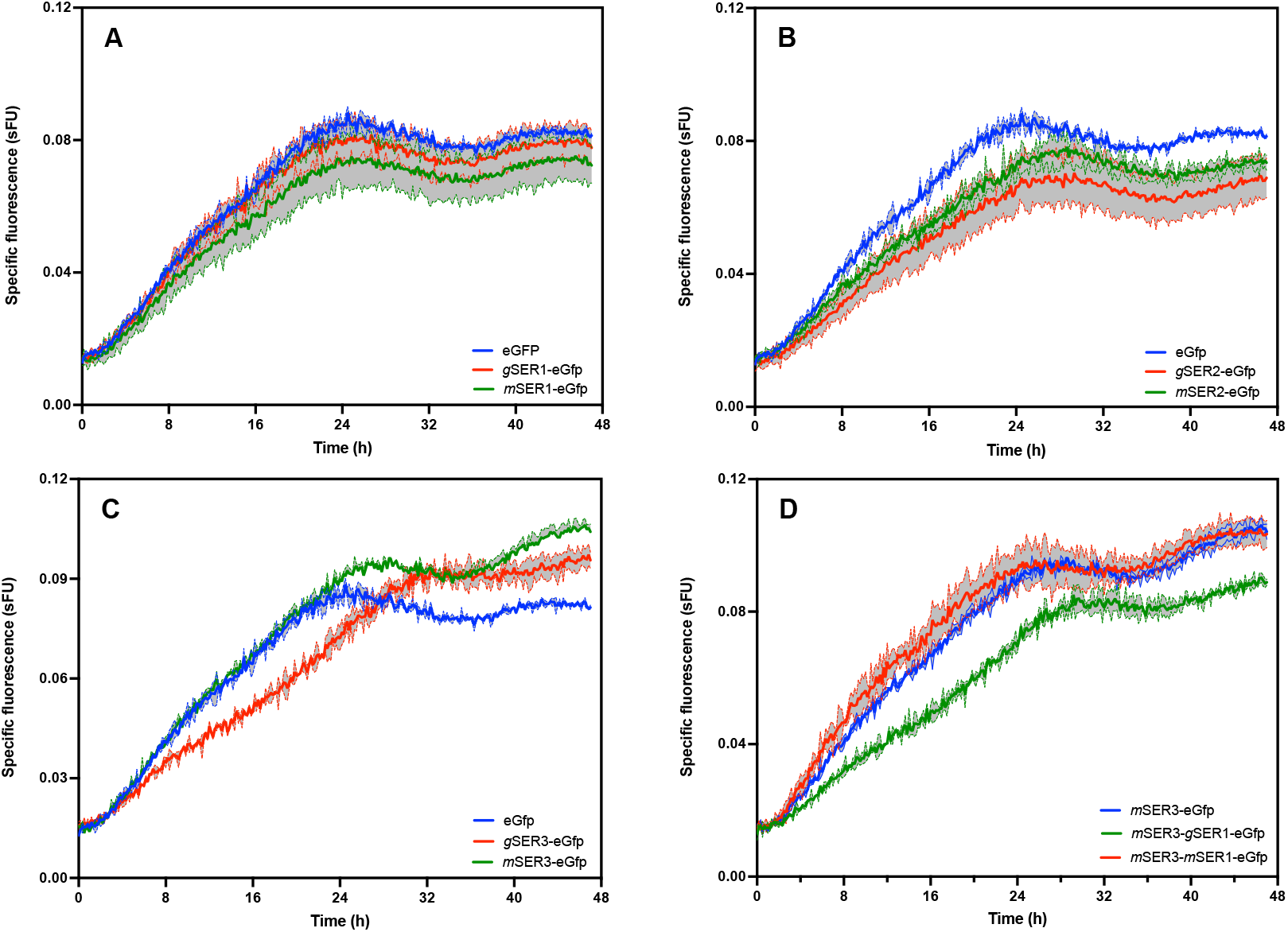
Specific eGFP fluorescence for eGfp strain (FdhKO pAOX1-eGFP) expressing gene SER1 (panel A), SER2 (panel B), SER3 (panel C) and SER1-SER3 (panel D) under the control the pGAP (strains marked with *g*) and pMDH3 (strains marked with *m*) during growth in YNBS. Cells were grown in BioLector in biological triplicates. Specific fluorescence are means (coloured lines) and standard deviation (grey shades). sFU: Specific fluorescence unit.

As SER1 expression is down regulated by serine in *K. phaffii* (Delic et al., 2014), the gene was co-expressed in the *m*SER3-eGFP background (fdh1Δ, pAOX1-eGFP, pMDH3-SER3) using both pGAP and pMDH3 promoters. The resulting strains, namely *m*SER3-*g*SER1-eGFP (fdh1Δ, pAOX1-eGFP, pMDH3- SER3, pGAP-SER1) and *m*SER3-*m*SER1-eGFP (fdh1Δ, pAOX1-eGFP, pMDH3-SER3, pMDH3-SER1) were cultivated in microbioreactor under the same experimental conditions (YNBS medium). Compared to the parental *m*SER3-eGFP strain, the specific fluorescence signal decreased when SER1 was expressed under the pGAP promoter (i.e., in the *m*SER3-*g*SER1-eGFP strain), whereas no significant change was observed when the pMDH3-driven expression (Figure 2). These results indicate that overexpression of SER1 in the *m*SER3-eGFP background does not further enhance pAOX1 induction.

In industrial rProt production processes based on pAOX1 driven expression systems, glycerol is commonly used as a feedstock during the cell growth phase, while a pAOX1 non-repressive carbon source, such as sorbitol, is preferred during the rProt production stage (Niu et al., 2013; Carly et al., 2016; Ergün et al., 2022;). Biomass and eGFP fluorescence were therefore monitored for the eGfp (*fdh1Δ*, p*AOX1-eGFP*) and *m*SER3-eGfp (*fdh1Δ*, p*AOX1-eGFP*, p*MDH3-SER3*) strains grown in microbioreactors on glycerol and sorbitol at a 1:2 ratio (YNBGS4). Both strains exhibited diauxic growth, consuming glycerol within the first 8 hours of culture (data not shown) followed by a sorbitol consumption phase (Figure 3). Both strains grew similarly on both carbon source, indicating that overexpression of SER3 does not affect cell fitness. However, marked differences were observed between the two strains regarding the specific fluorescence levels and thus pAOX1 induction. Upon glycerol depletion, eGFP-specific fluorescence markedly increased for both strains. The strain overexpressing SER3 (i.e. *m*SER3-eGfp) exhibited a 1.5-fold increase in specific fluorescence on average between 18 h and 48 h compared to eGfp strain (0.106 vs. 0.072 sFU, respectively). The fluorescence level then tended to decrease slightly as the biomass value reached a plateau (i.e. after 40 h).

**Figure 3:**
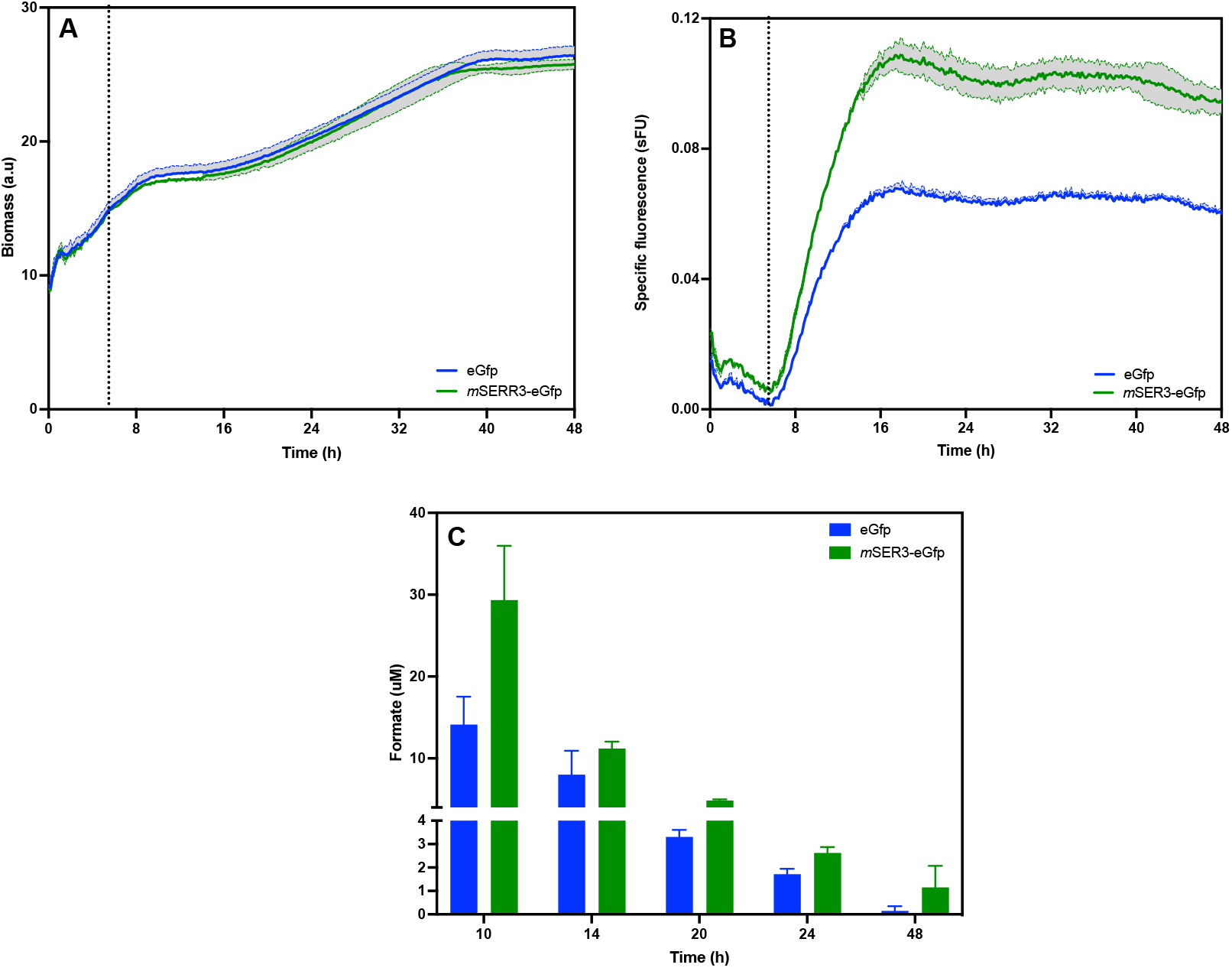
Biomass (panel A), specific eGFP (panel B) and formate concentration (Panel C) for eGfp (fdh1Δ, pAOX1-eGFP, blue color) and *m*SER3-eGfp (fdh1Δ, pAOX1-eGFP, pMDH3-SER3, green color) strains grown in YNB minimal media containing glycerol and sorbitol at a 1:2 ratio (YNBGS4). Cells were grown in a BioLector system, and data represent the mean and standard deviation of triplicate cultures. sFU: specific fluorescence unit; a.u: arbitrary units. μM: micromolar.

**Figure 4:**
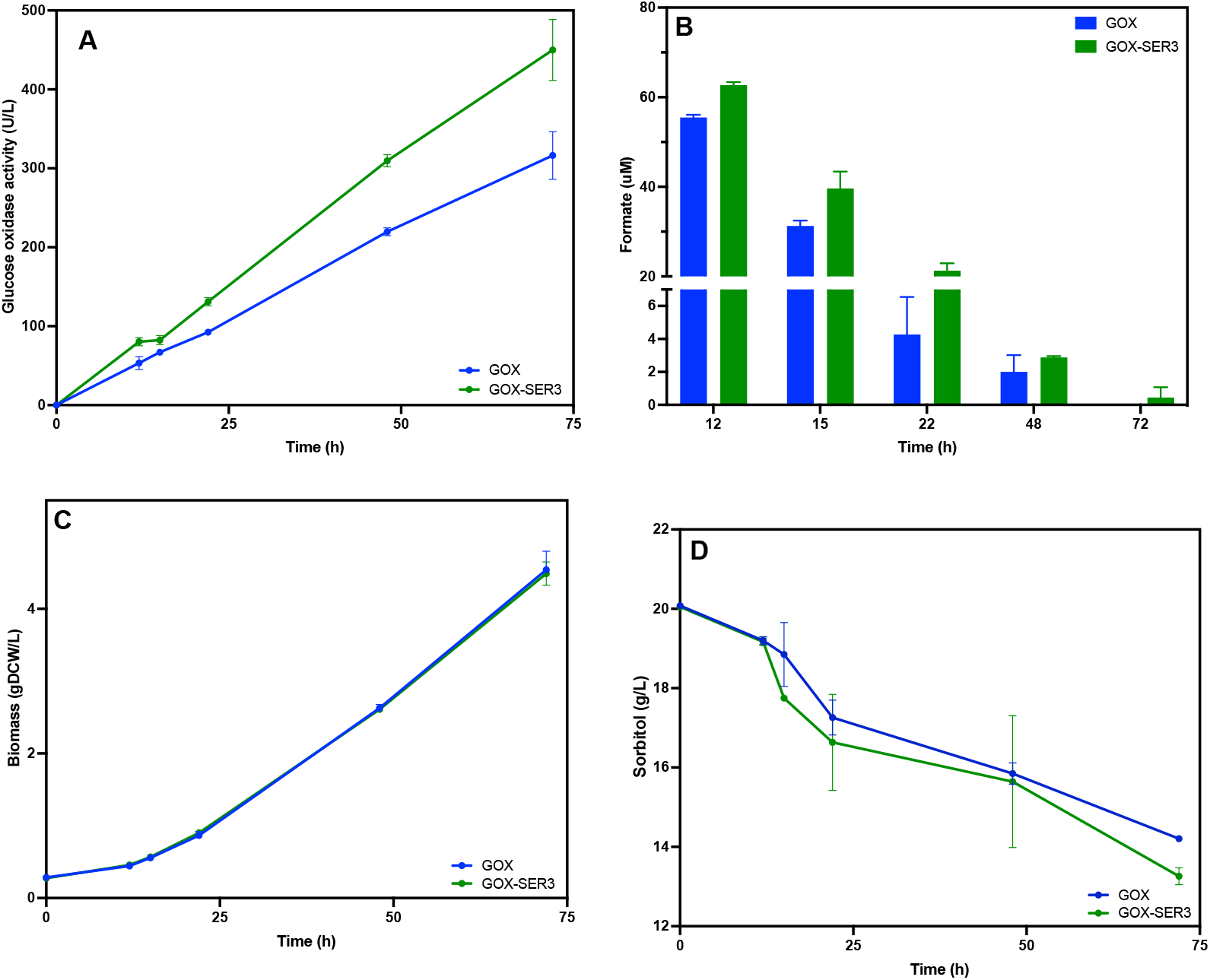
Glucose oxidase activity (panel A), formate concentration (panel B), biomass (panel C) and sorbitol concentration (panel D) during growth of GOX (blue colour) and GOX-SER3 (green colour) strains in sorbitol media. Data are the mean and standard deviation of triplicate cultures conducted in shake flasks. Glucose oxidase assays were performed in triplicates.

Formate concentration was also assessed at different time points. It increased for both strains after glycerol depletion to peak at 10 h of culture. Thereafter, the formate concentration gradually declined for both strains and became nearly undetectable by 48 h for the eGFP strain (Figure 3C). At any time points, formate content was consistently higher for the *m*SER3-eGFP strain compared to the eGFP strain, with the greatest difference observed at 10 h (2-fold, 29 μM vs 14 μM, respectively).

### Enhancing serine metabolism in FdhKO strains increases the synthesis of secreted protein in bioprocess conditions

The FdhKO pMDH3-SER3 strain (hereafter *m*SER3) was further considered for its ability to produce a secretory protein, namely glucose oxidase (Gox) from *A. niger*. Gox is an industrially relevant enzyme widely used in food processing, biosensing, and biocatalysis (Bankar et al., 2009). It is also a well-established model for secretory protein (Ahmad et al., 2014; Çelik & Çalik, 2012; Yu et al., 2020). The corresponding gene fused with the signal peptide from the α-mating factor from *S. cerevisiae* was cloned under the control of the pAOX1 promoter and expressed in a FdhKO and in the *m*SER3 strain. The resulting GOX (fdh1Δ, pAOX1-αMF-GOX) and GOX-SER3 (fdh1Δ, pAOX1-αMF-GOX, pMDH3-SER3) strains were grown on sorbitol (YNBS) in shake flasks. Cell growth, glucose oxidase activity and formate concentration were measured at various time points. After 72 h of cultivation, the GOX-SER3 strain exhibited a 1.4-fold increased Gox activity compared to the GOX strain, with respective values of 450 U/L and 316 U/L. Both strains showed similar growth kinetics (Fig. 3). Similarly to the eGFP strains, formate also accumulated for the GOX and GOX-SER3 strains (Fig. 3), and in a higher extend for the SER3-overexpressing strain. Specifically, formate accumulation peaked at 12 h of culture, reaching maximum values of 62 μM and 55 μM for GOX and GOX-SER3, respectively. Thereafter, formate concentration gradually declined in both strains and were nearly undetectable after 72 h, despite continued growth and with more than 70 % of the initial sorbitol still present in the culture medium (Figure 3). These results for a secreted proteins production are consistent with the trend observed with the eGFP strains (i.e. eGfp and *m*SER3-eGfp strains, Figure 2).

The GOX and GOX-SER3 strains were also cultivated in stirred-thank bioreactor under controlled conditions in medium containing glycerol and sorbitol at a 1:2 ratio (YNBGS). Cell growth, Gox specific activity, dissolved oxygen, and carbon source concentrations were monitored over 62 h. Both strains exhibited a diauxic growth, consuming glycerol the first 15h, followed by sorbitol during the subsequent 35 h. Glycerol depletion can be visualized by the sharp increase of dissolved oxygen at 15 h (Figure 5C). At this point, glycerol was no longer detectable in the culture supernatant, and sorbitol consumption has not yet begun (Figure 5B). During the sorbitol consumption phase, the dissolved oxygen levels remained around 85 % of saturation to finally increased to 100 % due to sorbitol depletion in the medium (Figure 5C). Throughout the entire cultivation period, oxygen was never a limiting substrate for cell metabolism. During the sorbitol consumption phase, both strains exhibited continuous growth at an average specific growth rate of 0.016 h^−1^, with a sorbitol uptake rate of 0.025 g·gDCW^−1^·h^−1^.

**Figure 5:**
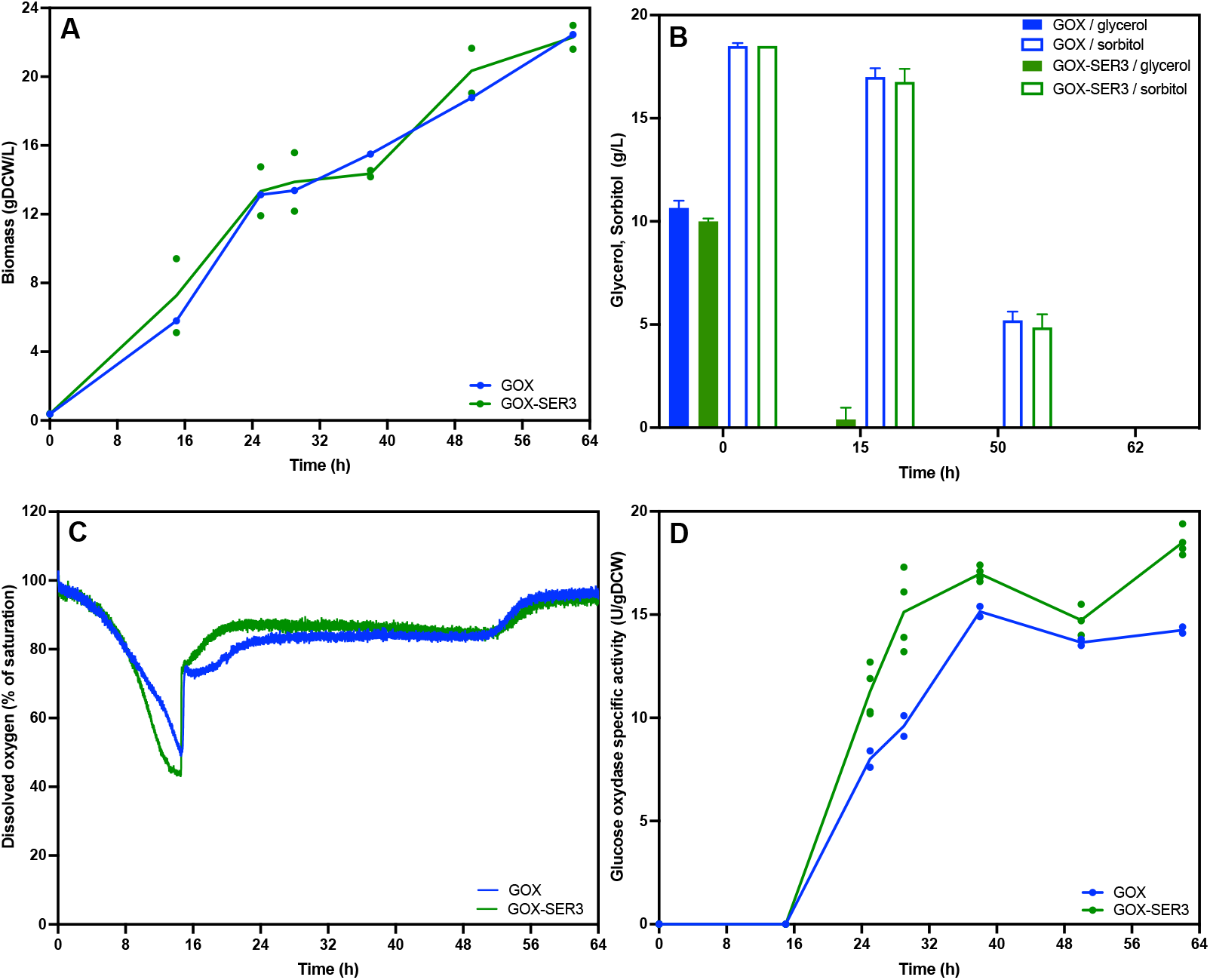
Biomass (panel A), glycerol (filled histograms) and sorbitol (empty histograms) concentrations (panel B), dissolved oxygen (panel C) glucose oxidase specific activity (panel D) during growth of GOX (blue colour) and GOX-SER3 (green colour) strains in stirred thank bioreactor in glycerol-sorbitol media (YNBGS). In panel A and D, the dots represent all the measurements while the lines connect the mean values.

No Gox activity was detected in the culture supernatant during the glycerol consumption phase, confirming that the pAOX1 promoter is tightly repressed by glycerol. Upon glycerol depletion, Gox specific activity (normalized to biomass) increased progressively in both strains, reaching a maximum after 40 hours and remaining stable until the end of the culture (62 hours). During growth on sorbitol, the Gox specific activity in the GOX-SER3 strain was 1.3-fold higher than that observed in the GOX strain (Figure 5D).

## Conclusion

We previously reported that disruption of the formate dehydrogenase gene in *K. phaffii* enables AOX1 promoter-based expression systems to function in a self-inducible manner, eliminating the need for methanol as an inducer. In this study, we demonstrated that overexpression of the gene encoding 3-phosphoglycerate dehydrogenase in a formate dehydrogenase-deficient *K. phaffii* strain significantly enhances *pAOX1* promoter induction under derepressed culture conditions. This strategy further improves rProt productivity, as exemplified by the enhanced secretion of Gox under process conditions.

## Author contribution

Rocio Cozmar: Strain construction, data acquisition; formal analysis writing the original draft; Patrick Fickers: Conceptualization; methodology, funding acquisition, supervision, editing the original draft; Julio Berrios: funding acquisition.

## Acknowledgements

The authors wish to thank Samuel Telek for his help with the bioreactor culture and Cristina Bustos for sharing some of the strains used in this study.

## Data availability statement

The data that support the findings of this study are available from the corresponding author upon reasonable request.

