## Supplementary figures and tables for "Engineering serine metabolism to enhance AOX1 promoter self-induction in a formate dehydrogenase-deficient *Komagataella phaffii*"

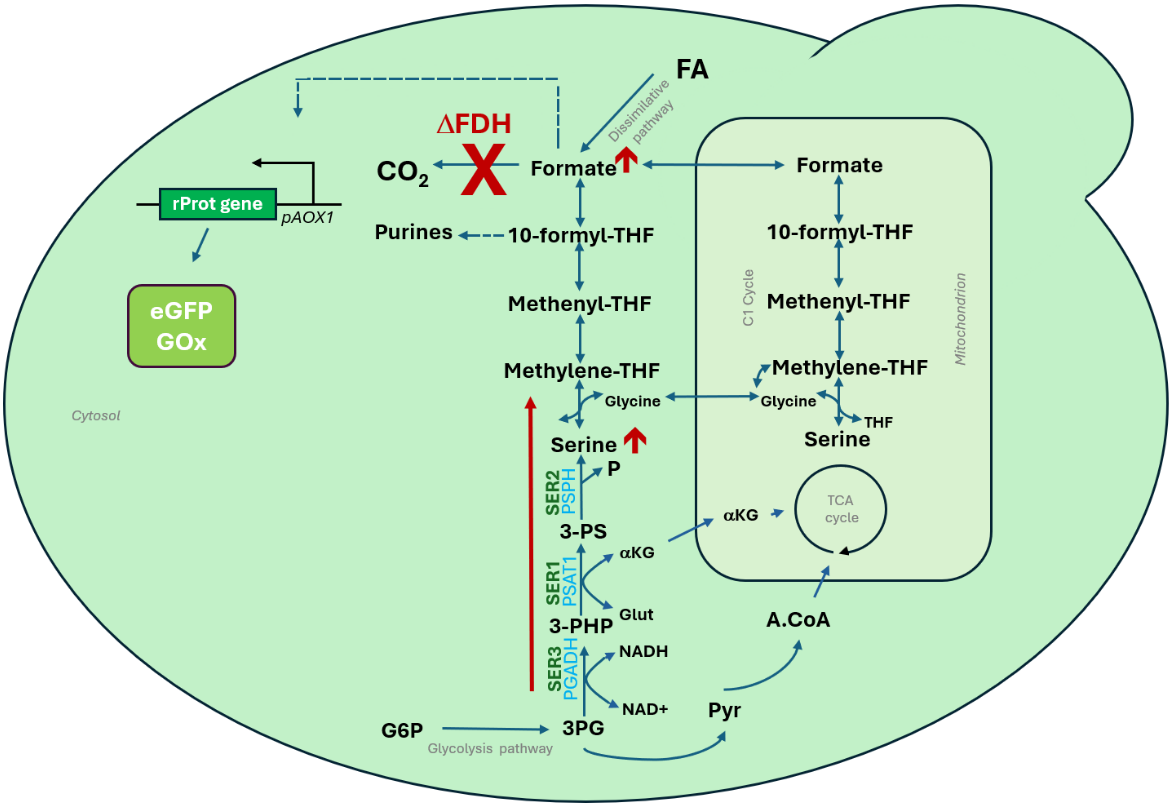


**Figure S1:** Schematic diagram illustrating the targeting serine biosynthesis pathway in a formate dehydrogenase-deficient *K. phaffii*. Formate from THF-C1 metabolism induces the p*AOX1* promoter.

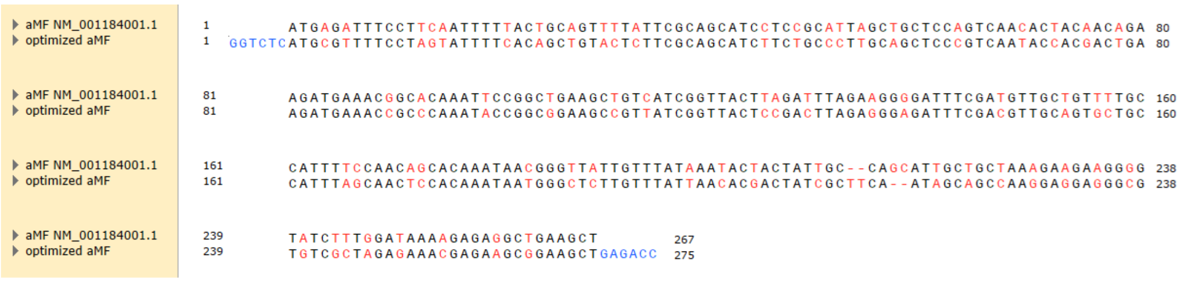

**Figure S2:** Codon optimized sequence of the alfa mating factor (αMF) from *Saccharomyces cerevisiae* (NM_001184001.1). Modified codons are in red, BsaI sequence is blue. Optimization was made using SnapGene software

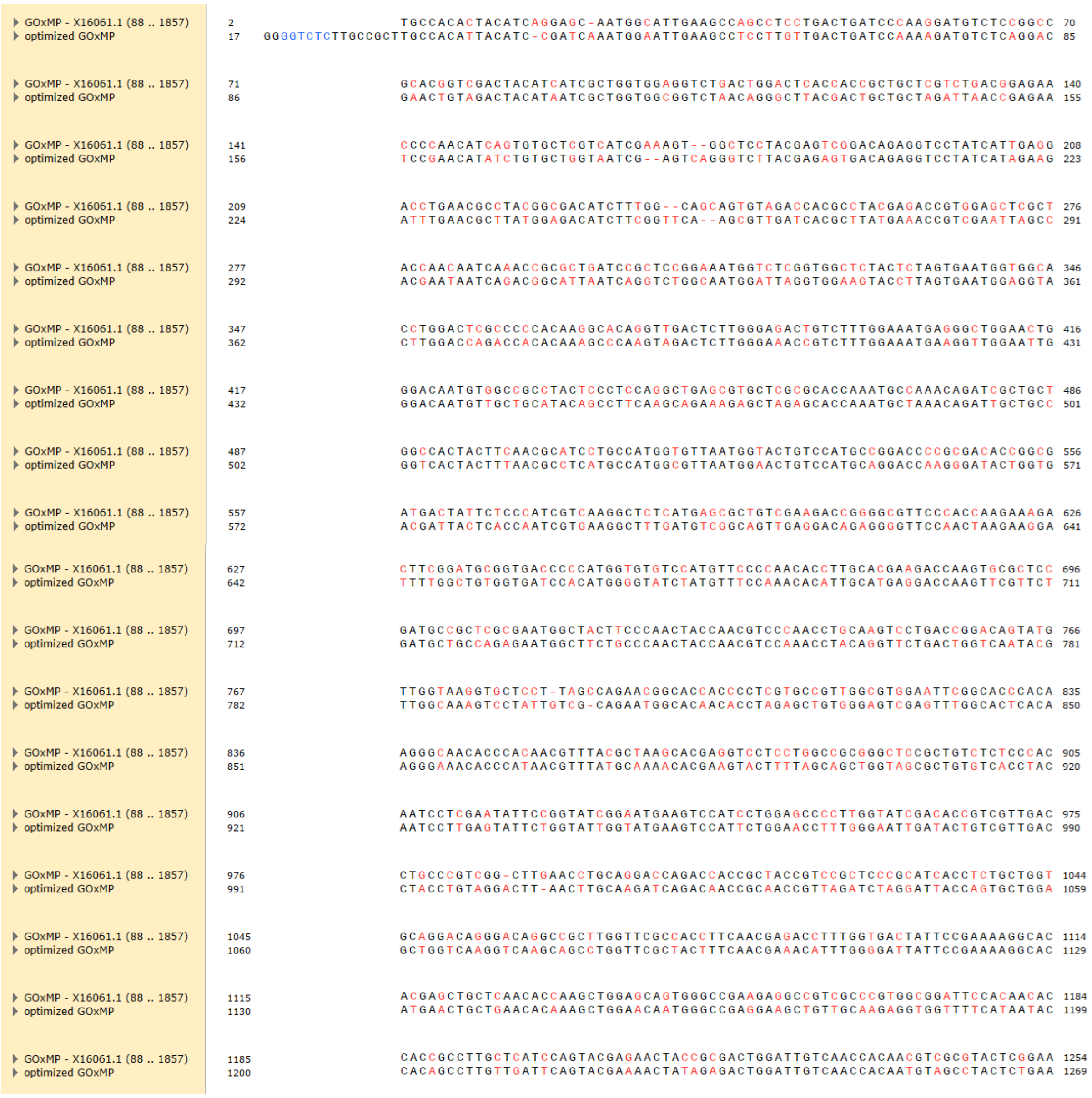

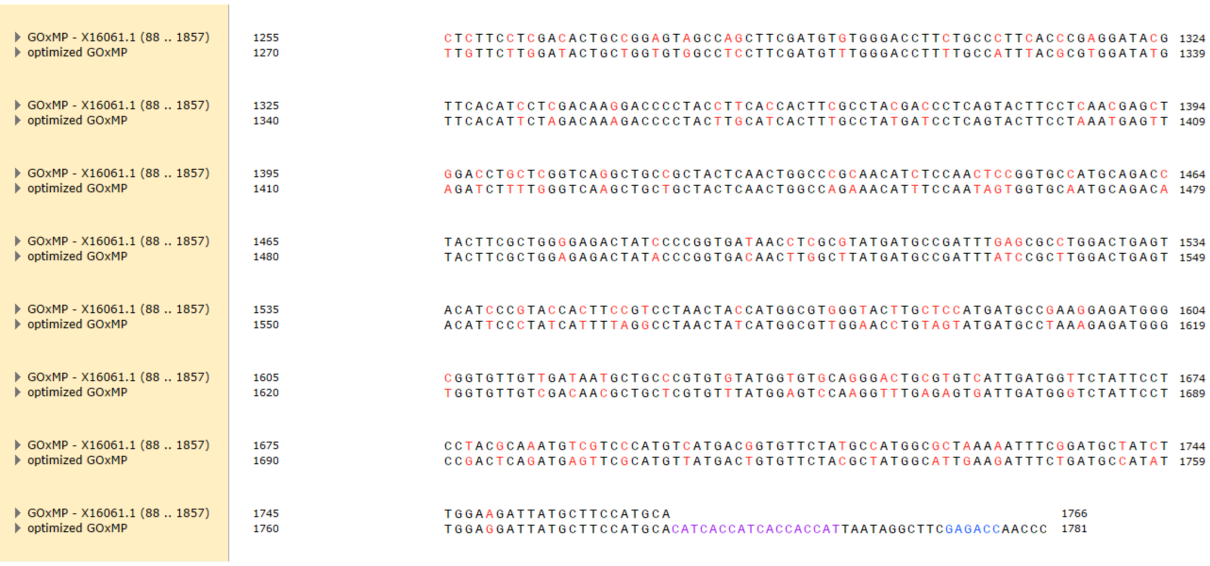


**Figure S3:** Codon optimized sequence of the intron less glucose oxidase gene from *Aspergillus niger* (x60601.1). Modified codons are in red. BsaI sequence is blue. Sequence is purple correspond to a histidine tag fuse to the Gox protein for purification. Optimization was made using SnapGene software.

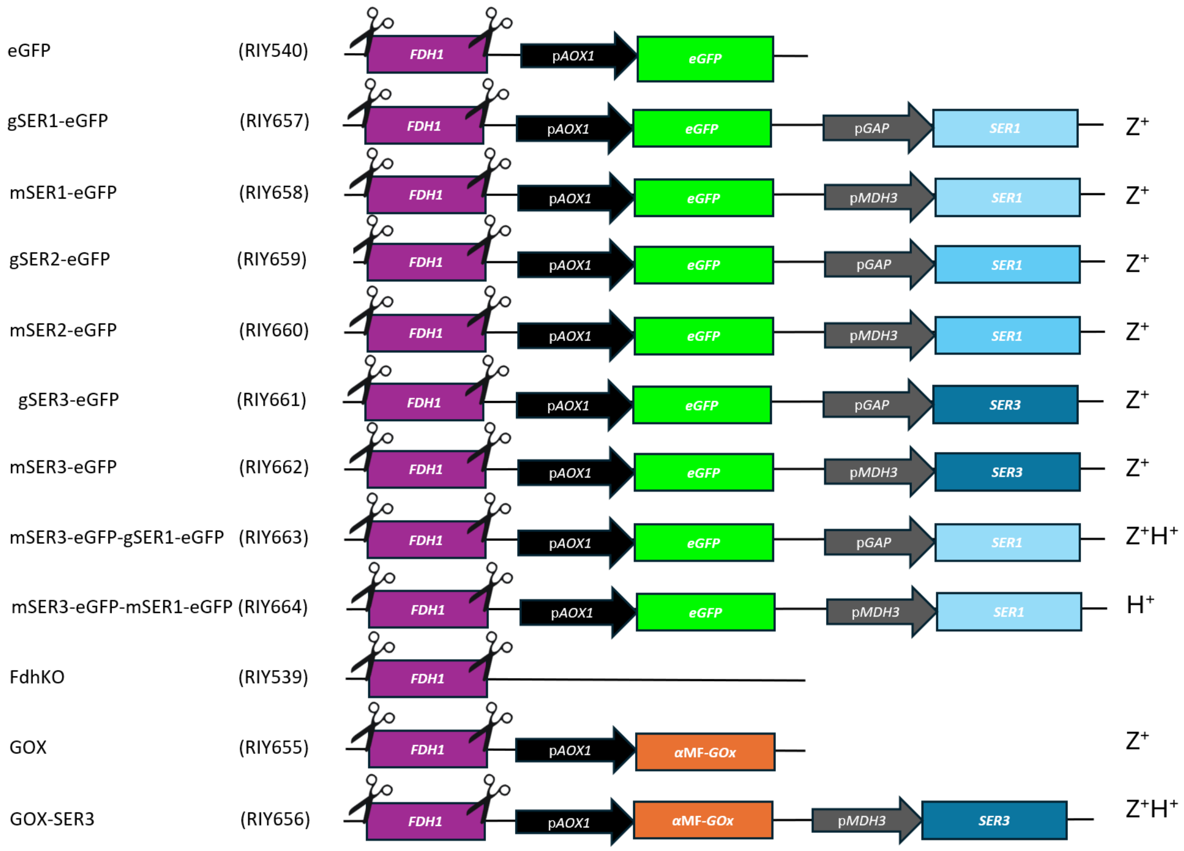


**Figure S4:** Schematic representation of the strain genotype used in the study.

**Table S1.** *Escherichia coli* strains used in this study.

| Strain name (plasmid) | Plasmid - genotype | Source/Reference |
|---|---|---|
| A2 | BB1_23 | (Prielhofer *et al.*, 2017) |
| D12 | BB3aZ_14 | (Prielhofer *et al.*, 2017) |
| E1 | BB3eH_14 | (Prielhofer *et al.*, 2017) |
| E8 | BB3rN_14 | (Prielhofer *et al.*, 2017) |
| A4 | BB1_12_p*GAP* | (Prielhofer *et al.*, 2017) |
| A9 | BB1_12_p*MDH3* | (Prielhofer *et al.*, 2017) |
| B8 | BB1_12_p*AOX1* | (Prielhofer *et al.*, 2017) |
| C1 | BB1_34_*ScCYC1*tt | (Prielhofer *et al.*, 2017) |
| RIE457 (RIP457) | pUC57-*aMF* (codon optimized) | This work |
| RIE485 (RIP485) | pGEMT-*GOX* ((codon optimized) | This work |
| RIE504 (RIP504) | A2_BB1_23_*aMFGOX* | This work |
| RIY508 (RIP508) | TOPO_*SER1* | This work |
| RIY509 (RIP509) | TOPO_*SER2* | This work |
| RIY510 (RIP510) | TOPO_*SER3* | This work |
| RIY511 (RIP511) | BB3aZ_14-p*GAP-SER1-SsCy1*tt | This work |
| RIE512 (RIP512) | BB3aZ_14-p*MDH3-SER1-SsCy1*tt | This work |
| RIE513 (RIP513) | BB3aZ_14-p*GAP- SER2-SsCy1*tt | This work |
| RIE514 (RIP514) | BB3aZ_14-p*MDH3-SER2-SsCy1*tt | This work |
| RIE515 (RIP515) | BB3aZ_14-p*GAP- SER3-SsCy1*tt | This work |
| RIE516 (RIP516) | BB3aZ_14-p*MDH3-SER3-SsCy1*tt | This work |
| RIE517 (RIP517) | BB3eH_14-p*GAP- SER1-SsCy1*tt | This work |
| RIE518 (RIP518) | BB3eH_14-p*MDH3-SER1-SsCy1*tt | This work |
| RIE519 (RIP519) | BB3rN_14-p*MDH3-SER3-SsCy1*tt | This work |
| RIE507 (RIP507) | BB3aZ_14-p*AOX1*- aMFGOX-*SsCy1*tt | This work |
| RIE396 (RIP396) | pKTAC-Cre | (Bustos et al., 2024) |

**Table S2**: Primers used in this study

| Name | Sequence 5´to 3´ |
|---|---|
| M13-Fw | gtaaaacgacggccagt |
| M13-Rv | aacagctatgaccatg |
| SER1-Fw | atggcaaaaactttggaaagagaa |
| SER1-Rv | tcaagaatgagcttcagcaa |
| SER2-Fw | atgagttatcgtttaactgc |
| SER2-Rv | ctattcgttaagaaatccct |
| SER3-Fw | atggcttctccccaagatat |
| SER3-Rv | ctagtatagcagacgggtga |
| 1.pGAP.SER1-Rv | **caaagtttttgccatggtgttttgatag**ttgttcaattgatt |
| 2.pGAP.SER1-Fw | **ctatcaaaacaccatggcaaaaactttg**gaaagagaagaacca |
| 1.pMDH3.SER1-Rv | **caaagtttttgccatgttgttattgttc**gttggttgtaactctaaagtct |
| 2.pMDH3.SER1-Fw | **gaacaataacaacatggcaaaaactttg**gaaagagaagaacca |
| 3.SER1.C1-Rv | **aaaaggggcctgaagctcaagaatgagcttc**agcaaattcgatg |
| 4.SER1.C1-Fw | **gaagctcattcttgagcttcaggccccttttt**cctttgtcgatatca |
| 1.pGAP.SER2-Rv | **taaacgataactcatggtgttttgatag**ttgttcaattgatt |

| | |
|---|---|
| 2.pGAP.SER2-Fw | **ctatcaaaacaccatgagttatcgttta**actgctatttcaaag |
| 1.pMDH3.SER2-Rv | **taaacgataactcatgttgttattgttc**gttggttgtaactctaaagtct |
| 2.pMDH3.SER2-Fw | **gaacaataacaacatgagttatcgttta**actgctatttcaaag |
| 3.SER2.C1-Rv | **aaaaggggcctgaagctctattcgttaaga**aatccctcaatttcg |
| 4.SER2.C1-Fw | **cttaacgaatagagcttcaggccccttttc**ctttgtcgatatcat |
| 1.pGAP.SER3-Rv | **tggggagaagccatggtgttttgatag**ttgttcaattgatt |
| 2.pGAP.SER3-Fw | **ctatcaaaacaccatggcttctcccca**agatattaagcagtta |
| 1.pMDH3.SER3-Rv | **ttggggagaagccatgttgttattgttc**gttggttgtaactctaaagtct |
| 2.pMDH3.SER3-Fw | **gaacaataacaacatggcttctccccaa**gatattaagcagtta |
| 3.SER3.C1-Rv | **aaaaggggcctgaagctctagtatagcaga**cgggtga |
| 4.SER3.C1-Fw | **tctgctatactagagcttcaggcccctttt**cctttgtcgatatcat |
| 4.BB3rN14.C1-Rv | **cattatacgaagttatggcatgccggagcg**agcgagcttgcaaattaaagcc |
| 1.BB3rN14pMDH3-Fw | **cagggcggggtttttcgcgatcggagg**tagcttgggtaggacttgacaagt |
| 4.BB3eH14.C1-Rv | **attctgggcctccatgtccatgccggagcg**agcgagcttgcaaattaaagcc |
| 2.pGAP-Fw | **gggcggggtttttcgcgatcggag**gatcctttttgtagaaatgtctt |
| 2.pMDH3-Fw | **gggcggggtttttcgcgatcggag**tagcttgggtaggacttgacaagt |
| 4.CYC1-Rv | **tcgacctgcagcgtaccatgccggagcg**agcttgcaaattaaagccttc |
| CkBB3aZ14-Fw | tgacagggcggggtttttcgcgatcggag |
| CkBB3aZ14-Rv | tcgacctgcagcgtaccatgccggagcg |
| 1.BB3rN14-Rv | ctccgatcgcgaaaaaccccgccctgtca |
| 4.BB3rN14-Fw | cgctccggcatgccataacttcgtataat |
| 1BB3aZ14-Rv | ctccgatcgcgaaaaaccccgccctgtcaggggcggggtttttg |
| 4.BB3eH14-Fw | cgctccggcatggacatggaggcccagaataccctccttgaca |

Overlap sequence for hifi assembly is in bold

**Table S3**: Enzymes involved in serine biosynthesis pathway in GS115 strain.

| Enzyme | Gene name | uniProtID GS115 | Gene ID |
|---|---|---|---|
| PGADH | SER3 | C4R1C8_KOMPG | PAS_chr2-1_0657 |
| PSAT1 | SER1 | F2QSD3_KOMPC | PAS_chr3_0566 |
| PSPH | SER2 | C4R7E7_KOMPG | PAS_chr4_0285 |
